# Genetic determinism of spontaneous masculinisation in XX female rainbow trout: new insights using medium throughput genotyping and whole-genome sequencing

**DOI:** 10.1101/2020.04.28.057240

**Authors:** Clémence Fraslin, Florence Phocas, Anastasia Bestin, Mathieu Charles, Maria Bernard, Francine Krieg, Nicolas Dechamp, Céline Ciobotaru, Chris Hozé, Florent Petitprez, Marine Milhes, Jérôme Lluch, Olivier Bouchez, Charles Poncet, Philippe Hocdé, Pierrick Haffray, Yann Guiguen, Edwige Quillet

## Abstract

Rainbow trout has a male heterogametic (XY) sex determination system controlled by a major sex-determining gene, *sdY*. Unexpectedly, a few phenotypically masculinised fish are regularly observed in all-female farmed trout stocks. To better understand the genetic determinism underlying spontaneous maleness in XX-rainbow trout, we recorded the phenotypic sex of 20,210 XX-rainbow trout from a French farm at 10 and 15 months post-hatching. The masculinisation rate was 1.45%. We performed two genome-wide association studies (GWAS) using both medium-throughput genotyping (31,811 SNPs) and whole-genome sequencing (WGS, 8.7 million SNPs) on a subsample of 1,139 individuals classified as females, intersex or males. The genomic heritability of maleness ranged between 0.48 and 0.62 depending on the method and number of SNPs used for estimation. At the 31K SNPs level, we detected four QTLs on three chromosomes (Omy1, Omy12 and Omy20). Using WGS information, we narrowed down the positions of the two QTLs detected on Omy1 to 96 kb and 347 kb respectively, with the second QTL explaining up to 14% of the total genetic variance of maleness. Within this QTL, we detected three putative candidate genes, *fgfa8, cyp17a1* and an uncharacterised protein (LOC110527930), which might be involved in spontaneous maleness of XX-female rainbow trout.

## Introduction

The discovery of factors underlying sex determination in fish is a challenge for fundamental biological and evolutionary perspectives and for aquaculture purposes as in some species, managing sex ratios of farmed stocks is essential for the production efficiency. Unlike birds and mammals which have highly conserved and simple heterogametic genetic sex determination systems, teleost fish species exhibit an amazing diversity of sex determination systems, with different master sex determining genes as well as many minor genetic determinants interacting with environmental effects (in particular temperature) and epigenetic mechanisms^1–3^. Among teleost fish, salmonids are known as strict gonochoristics with the individual sex determined at fertilization and remaining the same throughout their live^1^. In rainbow trout (*Oncorhynchus mykiss*), the genetic determinism of sex has first been described as a male heterogametic mechanism (males XY, females XX) based on observations in progenies of hormonally sex-reversed individuals^4,5^ and the absence of male in gynogenetic offspring^6^. The master gene controlling sex determination in salmonids^7,8^ is an unusual sex determining gene called *sdY* (sexually dimorphic Y chromosome) that evolved from the duplication of the *irf9* immune related gene. *sdY* is expressed during early differentiation in the testis where it interacts with the conserved female differentiation factor Foxl2^9^ to prevent *cyp19a1a* up-regulation and the subsequent oestrogen production needed for ovarian differentiation^10^. However, despite a strict sex-linkage of the *sdY* locus in many salmonid species^8^, spontaneous masculinisation of XX females have been reported in rainbow trout, first in experimental groups^11^, but also in commercial populations (unpublished data). These spontaneous masculinisation and their transmission across generations have been characterized in gynogenetic families, where the role of minor genetic factors acting in addition to the major *sdY* sex determination system has been suspected^11,12^. Later, Guyomard et al.^13^ (2014) detected QTLs associated to masculinisation in two doubled haploid gynogenetic trout families. In addition to this genetic control, a few studies have also shown that high temperature treatments applied before the sexual differentiation and at the period of thermosensitivity can enhance the frequency of maleness in rainbow trout^14,15^, as shown in some other species including for instance channel catfish (*Ictalurus punctatus*), Nile tilapia (*Oreochromis niloticus*) and seabass (*Dicentrarchus labrax*)^16,17^. Interestingly, in most of the studies in trout, intermediate phenotypes (intersex individuals with only partial masculinisation of gonads) were recorded.

In rainbow trout farming, the production of all-female populations is advantageous due to their late sexual maturation (at 2 years-old) compared to males (about 1 year of age), because maturing males exhibit a reduced growth and flesh quality (reduction of muscle lipid content and discoloration) and are more sensitive to fungal saprolegnosis. In all-female stocks, reproduction is performed by mating standard XX-females with sex-reversed XX-males. Sex reversal is obtained by feeding young trout fry with a diet containing a masculinising hormone, the 17α-methyltestosterone before the sexual differentiation period^18^. The administration of hormones to obtained sex-reversed males is carried out under strict veterinary controls according to the European directive (99/22/CE, 29 April), and treated animals are euthanized after reproduction and discarded from the food chain.

Therefore, there is a dual interest in investigating factors that govern the spontaneous maleness in trout. Depending on the nature of those factors, they could be exploited either to reduce the occurrence of undesirable males observed in all-females populations or to produce sex-reversed males without using hormones, an asset for more sustainable aquaculture.

To further investigate the genetic architecture of this spontaneous maleness, we performed a Genome-Wide Association Study (GWAS) in a French commercial rainbow trout population, in which spontaneous maleness has been repeatedly reported. Genotype information obtained with a medium-density trout genotyping array and whole genome sequence (WGS) information were combined to accurately map QTL and detect candidate genes responsible for spontaneous maleness.

## Methods

### Fish rearing

In June 2017, at the French trout farm « *Les Fils de Charles Murgat* » (Beaufort, France), 2.5 years-old breeders were reproduced after photoperiodic induction of maturation. In total, 50 dams (XX) and 50 sex-reversed sire (XX) were mated according to 5 full-factorial mating designs (10 dams x 10 sires per factorial) to produce all XX-eggs. Fertilized eggs were separated into two batches corresponding to two temperature treatments (25,000 and 45,000 eyed eggs per batch) and incubated at 10.5°C until hatching. Hatching rate was 64% and 58% in the two batches, which is in the usual range when using progenitors whose spawning has been shifted by photo-period. At the end of yolk resorption (35 days post-fertilisation, dpf), larvae were kept at 12°C in the first group (control) while in the second group, the water temperature was increased by 1°C/day until it reached 18°C. The water temperature was then maintained constant for 1134 degree-days covering the expected window of gonad differentiation^12^. It was then decreased by 1°C/day during 6 days to reach the same temperature as in the control group (12°C) and both groups were then reared between 12 to 14°C. At 84 dpf, 13,000 fish from each temperature group were randomly sampled and kept in the experiment. In order to prevent too much size heterogeneity within the groups and any risk of biased sampling at time of phenotyping, each group was temporarily split (at 214 dpf) into two subgroups according to the fish size and kept separated for 37 days, allowing the smaller fish to catch up in size with the bigger fish. Fish were then regrouped until sex was recorded. During the whole experiment, fish were fed a commercial diet according standard recommendations. As part of the standard breeding practices in a commercial broodstock, the handling of fish was not subject to oversight by an institutional ethic committee.

### Phenotypic sex recording

In order to determine the phenotypic sex of the XX-offspring, 13,241 fish from both control and 18°C groups (6,560 and 6,681 fish, respectively) were sexed at 10 month-post fertilisation (mpf) and the remaining 6,969 fish were sexed 5 months later (3,456 and 3,513 fish, respectively). The overall mean body weight of fish was respectively 234g at 10 mpf and around 860g at 15 mpf. Fish were euthanized by electro-narcosis, according to standard procedures for commercial trout farming and both gonads were visually examined to determine sex. When necessary, visual observation was confirmed by observation of the gonads under a binocular magnifier (1,079 fish) and, in some cases, with a histological control (23 fish).

Fish were distributed in four sex classes (no matter the degree of maturation) as followed:

- Female, for fish with two ovaries observed and no visible sign of testis area (See Supplementary Figure S1, panel a for an example).
- Intersex, for fish with either one entirely female gonad and one entirely male gonad, or with both male and female areas in at least one of the gonads (the other gonad may be male, female, intersex or undetermined) (n = 132). The presence of ovarian lamellae was the criterion used to declare a female area in a gonad, whether oocytes were present or not (See Supplementary Figure S1, panel b for an example)..
- Male, for fish with two testis observed, or for five fish, one testis only (the other gonad being undifferentiated) and no visible sign of any ovary area (n = 162) (See Supplementary Figure S1, panel c for an example)..

Fish with undifferentiated gonads that could not be sexed after binocular or histological observation were classified as undetermined and removed from the analysis (n = 84).

All intersex trout (n=132), all males (n=162) as well as 858 females with well-developed ovaries were kept for QTL detection (563 from the control group, at both 10 and 15 mpf, and 295 from the 18°C group at 15 mpf). Sex was recorded as a categorical variable with three levels as follow: sex = 1 for females, sex = 2 for intersex, sex = 3 for males.

### Genotyping and sequencing

Pieces of caudal fin sampled from those 1,152 fish were sent to Gentyane genotyping platform (INRAE, Clermont-Ferrand, France) for DNA extraction using the DNAdavance kit from Beckman Coulter following manufacturer instructions. Genotyping was performed with the Axiom™ Trout Genotyping Array from Thermofisher^19^ that contains 57,501 SNPs.

Quality controls of genotyped SNPs were performed as described in D’Ambrosio *et al*.^20^ in particular to remove SNPs with probe polymorphism and multiple locations on the genome. In addition, only the 31,811 SNPs with a call rate higher than 0.97, a test of deviation from Hardy-Weinberg equilibrium with a p-value > 0.0001 and a minor allele frequency (MAF) higher than 0.05 were conserved for the analysis. Individuals with a call rate lower than 0.90 were removed from the genotype dataset (n=5). All missing genotypes were imputed using the FImpute software^21^ (version 2.2) in order to get the full 31,811 SNPs genotypes for the animals considered in the analysis.

Samples of caudal fin were also collected from a set of 60 females (50 dams + 10 relatives) for DNA extraction and whole genome sequencing. DNA was extracted with a Promega Relia Prep dDNA tissue miniprep system (A2051) following manufacturer instructions. DNAseq was performed at the GeT-PlaGe core facility (INRAE, Toulouse, France, https://get.genotoul.fr/en/). DNA-seq were prepared according to Illumina’s protocols using the Illumina TruSeq Nano DNA HT Library Prep Kit. Briefly, DNA was fragmented by sonication, size selection was performed using SPB beads (kit beads) and adaptors were ligated to be sequenced. Library quality was assessed using Advanced Analytical Fragment Analyzer and libraries were quantified by QPCR using Kapa Library Quantification Kit. DNA sequencing was performed on an Illumina NovaSeq6000 using a paired-end read length of 2×150 bp with the Illumina NovaSeq6000 S4 Reagent Kits.

After sequence mapping on the reference genome assembly^22^ (bwa mem v0.7.17, GCA_002163495.1), 34,064,394 variants were obtained using a homemade pipeline (https://forgemia.inra.fr/bios4biol/workflows/tree/master/Snakemake/IMAGE_calling) with three variant calling tools (GATK v3.7.0, FreeBayes v1.2.0 and SAMtools mpileup v1.8.0). Quality controls and filtering were performed using vcftools^23^ (version 1.15). First, indels and SNPs on un-located contigs or mitochondrial DNA were removed. Only the 21,904,314 biallelic SNPs located on identified chromosomes were further considered in the analysis. Then, a filtering was performed on variant coverage, with the removal of variants with either less than 10X reads or more than 50X reads (considered as putative duplicated regions). Variants with more than 58 individuals being homozygous for either the reference or the alternative allele were removed. In other words, 14,478,077 variants with at least two individuals different from the 58 others (either heterozygous or homozygous for the alternative alleles) were kept. Finally, only the 8,784,147 variants with a MAF equal or above 10% were conserved for the GWAS analysis and imputation.

### Imputation to whole-genome-sequence

Imputation to whole-genome-sequence (WGS) of the 31K SNPs genotypes of the progeny was performed chromosome by chromosome using the latest version of FImpute software^21^ (version 3.0) based on the reference population constituted by the 60 sequenced females. After imputation, 8,765,613 SNPs with a MAF higher than 1% were kept for the WGS analysis. In order to reduce the variant dataset to produce a genomic relationship matrix, SNPs in linkage disequilibrium were filtered out with the indep-pairwise option of the PLINK software^24^ (version 1.09). The filtering was first performed on 50 bp sliding windows, r^2^ between each pair of SNPs in the window was calculated and one SNP for the pair was removed if the r^2^ was higher than 0.7 for a first filtering and higher than 0.3 for a second filtering performed on a 100 bp sliding window. The reduced dataset obtained was composed of 275,283 SNP.

### Genome Wide Association Studies

#### Models

Genome wide association studies (GWAS) were performed at the genotyping level on the 31K SNPs dataset and at the sequence level (WGS) on the 8.7 million SNPs dataset using two different approaches, a marker-by-marker analysis and a Bayesian Stochastic Search Variable Selection approach.

The first GWAS approach was performed with the GCTA software^25^ that performs a marker-by-marker analysis under a mixed linear model with a correction for data structure based on a genomic relationship matrix^26^ (GRM).

The model used in this first GWAS is described by the equation (1):

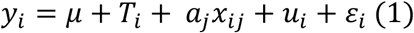

With y_i_ the observed phenotype for the i^th^ individual (sex in 3 levels), *μ* the overall mean in the population, T_i_ the fixed effect of the temperature treatment (2 levels), a_j_, the additive effect of the reference allele of the candidate SNP (j) to be tested as fixed effect for sex association and x_ij_ the reference allele count (0,1, or 2) for the SNP j for individual i, u_i_ is the random polygenic additive value of individual i and ε_i_ the residual effect for individual i. The vector of residual effects is normally and independently distributed 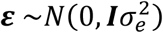 with σ^2^e the residual variance. The random vector of polygenic effects followed a normal distribution 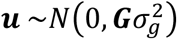 with 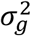 the estimated genetic variance and **G** a GRM constructed using only SNPs information. We first performed a GWAS on the 31K SNPs level with a GRM constructed using all 31K SNPs included in the model, this approach will be referred to as GCTA-chip (see Table 1). In order to refine the position and size of the QTL detected by the analysis at the 31K SNPs level, we performed similar GWAS using the WGS dataset. This GWAS at the sequence level was performed on the whole 8.7 million SNPs with a GRM constructed with 275K SNPs according to Yang *et al*.^26^. This GWAS will be referred to as GCTA-seq for the rest of this article (Table 1).

**Table 1.** Summary of GWAS approaches used to detect QTL associated with spontaneous maleness in XX-rainbow trout. π is the number of SNPs with a non-zero-effect included at each cycle k of the MCMC algorithm. GRM: Genomic Relationship Matrix

| Name | Software | Number of SNP in the analysis | Number of SNP in GRM | $\pi$ |
| --- | --- | --- | --- | --- |
| GCTA_chip | GCTA | 30,811 | 30,811 | - |
| GCTA_seq | GCTA | 8,765,613 | 275,283 | - |
| BCπ-chip | BESSiE | 30,811 | 275,283 | 1 % |
| BCπ-seq | BESSiE | 21,149 | 275,283 | 0.02 % |

The second GWAS approach was performed using the Bayesian Stochastic Search Variable Selection approach BayesCπ^27^ implemented in the BESSiE software^28^ (version 1.0). In this BayesCπ approach at each iteration, a proportion π of SNP is assumed to have a non-zero-effect on the trait, thus at each iteration the number of SNP effects to be estimated is lower than the number of phenotypic records. The statistical model used is defined according to equation 2:

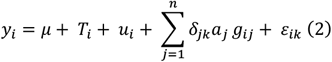

With y_i_ the observed phenotype the i^th^ individual (sex in 3 levels), μ the overall mean in the population, T_i_ the fixed effect of the temperature treatment (2 levels), u_i_, the additive polygenic effect for individual i, a_j_, is the additive effect of the reference allele for the j^th^ SNP with its genotype for individual i (g_ij_) coded as 0, 1 or 2 and n the number of SNP in the analysis; δ_jk_ is the indicator variable for the non-zero effect of the j^th^ SNP at the k^th^ iteration; *ε_ik_* the residual effect for the i^th^ individual at the k^th^ iteration. As in the previous model, the random vector of polygenic effects followed a normal distribution 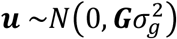 with 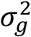 the estimated genetic variance and **G** the GRM constructed using information from the 275K SNPs. The vector of residual effects is normally and independently distributed 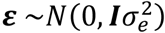 with σ^2^e the residual variance.

At each cycle k, the decision to include SNP j in the model depends on the indicator variable δ_jk_, with the effect (a_j_,) of the SNP j estimated if δ_jk_ is equal to 1, and not estimated if δ_jk_ is equal to 0. This indicator variable^27^ is sampled from a binomial distribution with a probability π that δ_jk_ is equal to 1 (the SNP as a non-zero-effect) and a probability 1-π that δ_jk_ is equal to 0.

As for the marker-by-marker approach, a first GWAS using the BayesCπ model was performed on the 31K SNPs level with determined parameters in order to have 1% of the marker included in the model at each cycle. This approach will be referred to as BCπ-chip (Table 1). To refine the localisation of QTL, the second GWAS at the WGS level was performed using only SNP located on a portion of one chromosome (selected using the results of the GCTA-seq analysis) with the effect of all other SNPs (from that chromosomes and others) included in the polygenic effect calculated with the GRM estimated with the 275K SNPs. For this approach, which will be referred to as BCπ-seq (Table 1), the proportion π of SNPs to be included in the analysis at each cycle was constrained to 0.02% (i.e. 3 SNPs included at each iteration).

Both BayesCπ analyses were performed with a MCMC algorithm. A total of 500,000 iterations were used for the BCπ-chip and 1 million iterations were used for the BCπ-seq, with, for both analyses, a burn-in period of 50,000 cycles and results saved every 20 cycles. In order to control the reproducibility of our analyses, they were both run twice, with two different seeds for the random number generator to initialize MCMC algorithm. Convergence was assessed by visual inspection of trace plots of the SNP effects and variance components estimates for all different seeds.

### Estimation of genetic parameters

Variance components and heritability were estimated using the average information of restricted maximum likelihood analysis (AI-REML) implemented in GCTA software^25^ with the same model as equation (1). The two GRM build with 31K and 257K SNPs for GWAS were used separately to estimate genetic parameters at the genome and the sequence level.

The values obtained with the 31K SNPs GRM were used as prior for both BayesCπ models and genetic variance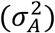 was calculated as the sum of the polygenic variance (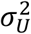, estimated by BESSiE) and the genomic variance 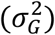 estimated by n SNPs calculated according to (3):

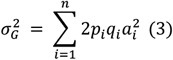

With p_i_ and q_i_ the allele frequencies for the i^th^ SNP and a_i_ the estimated additive effect of the i^th^ SNP.

Genomic heritability (h^2^_g_) was estimated as 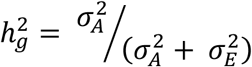 with 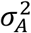 the estimated genetic variance and σ^2^_E_ the residual variance.

### QTL definition

For the GCTA-chip and GCTA-seq analysis, we determined chromosome-wide suggestive and genome-wide significance thresholds using a Bonferroni correction of α = 1% (genome-wide threshold = −log_10_(α/n); chromosome-wide threshold = −log_10_(α/[n/30]) with n the number of SNP in the analysis (30,811 or 8.7 millions). Only SNPs with −log_10_(P-value) over the chromosome-wide threshold for GCTA-chip and over the genome-wide threshold for GCTA-seq were considered. Approximate confidence intervals (CIs) for each QTL with a peak SNP value over the significance threshold were estimated using a drop-off limit^29^ of 1.5 unit of −log_10_(P-value) and a maximum distance of 1 Mb for the GCTA-chip and 200 kb for the GCTA-seq between two successive SNPs with −log_10_(P-value) over the drop-off limit starting from the position of the peak SNP.

In the GCTA-seq, when two successive QTLs were detected at a distance lower than 50 bp and with less than 250 kb between their peak SNPs, the CIs were cumulated in a credibility interval (See Supplementary Table S1 for detailed CIs).

For the Bayesian approaches, the degree of association between a SNP and the phenotype was assessed using the Bayes Factor^30^ (BF):

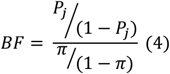

With P_j_ the probability of the SNP j having a zero effect and π the proportion of SNP having a non-zero effect on the trait (in our case π = 1% for BCπ-chip and π = 0.02% for BCπ-seq). BF was transformed into logBF computed as twice the natural logarithm in order to obtain values of the same usual range as the P-value, thus facilitating the visual appraisal of QTL and the comparison between methods^31^. To defined QTLs with the BayesCπ approaches, we defined two categories to classify the strength of the evidence in favour of a QTL^32^: evidence was consider as strong for 8 ≤ logBF < 10 and as very strong for logBF ≥ 10. Because the Bayes Factor is not a statistic test, confidence intervals cannot be derived, but credibility intervals can be built as defined in Michenet *et al*.^33^. Credibility intervals were determined using the threshold logBF ≥ 8 for defining a peak SNP showing evidence for a QTL in either a BCπ-chip or a BCπ-seq analysis. For the BCπ-chip approach, the credibility interval included every SNP with a logBF > 3 within a 1 Mb sliding window from the peak SNP. For the BCπ-seq approach, it included every SNP with a logBF > 5 within a 200 kb sliding window from the peak SNP.

The proportion of genetic variance explained by each SNP was derived from the Bayesian analyses and calculated according to (5):

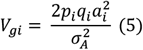

With p_i_ and q_i_ the allele frequencies for the i^th^ SNP, a_i_ the estimated additive effect of the i^th^ SNP and 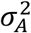 the total additive genetic variance (as defined earlier). To estimate the proportion of variance explained by a QTL, the proportions of genetic variance explained by all the SNPs within the credibility interval were cumulated.

### Candidate genes and SNP annotation

Candidate genes located within a reduced interval determined as the intersection of confidence and credibility intervals estimated using both GCTA-seq and BCπ-seq analysis were listed from the NCBI *Oncorhynchus mykiss* Annotation Release 100 (GCF_002163495.1).

Annotation of SNPs within those intervals was performed using the SNPEff software^34^ (version 4.3) with the NCBI *Oncorhynchus mykiss* Annotation Release 100 (GCA_002163495.1)^22^ as a reference. SNPs were then filtered according to their estimated putative impact, all SNPs with a modifier putative impact were filtered out and only SNP with a low, moderate, or high putative impact were conserved.

### Pedigree and dams’ genotypes

Genomic pedigree information’s were recovered using identity by descent (IBD) estimates from PLINK software^24^ (version 1.9) using information of the 31K SNPs. The percentage of masculinised fish in the genotyped offspring of each dam was calculated (see Supplementary Table S2). The 50 dams used in the mating scheme were named according to the proportion of masculinised fish in their offspring with the AA-dam having the higher proportion of masculinised offspring and the BX-dam having the lower proportion of masculinised offspring (Figure 1). Among the dams with at least 10 progeny recorded, the 22 dams with extreme progeny phenotypes were selected: the 11 dams (labelled from BH to BR) that had less than 8% of masculinised fish in their offspring and the 11 dams (labelled from AA to AK) that had more than 35% of masculinised offspring. Those 22 dams with extreme masculinisation rate in the offspring sample were selected for in-depth analysis of their genotypes at the QTL regions.

**Figure 1.**
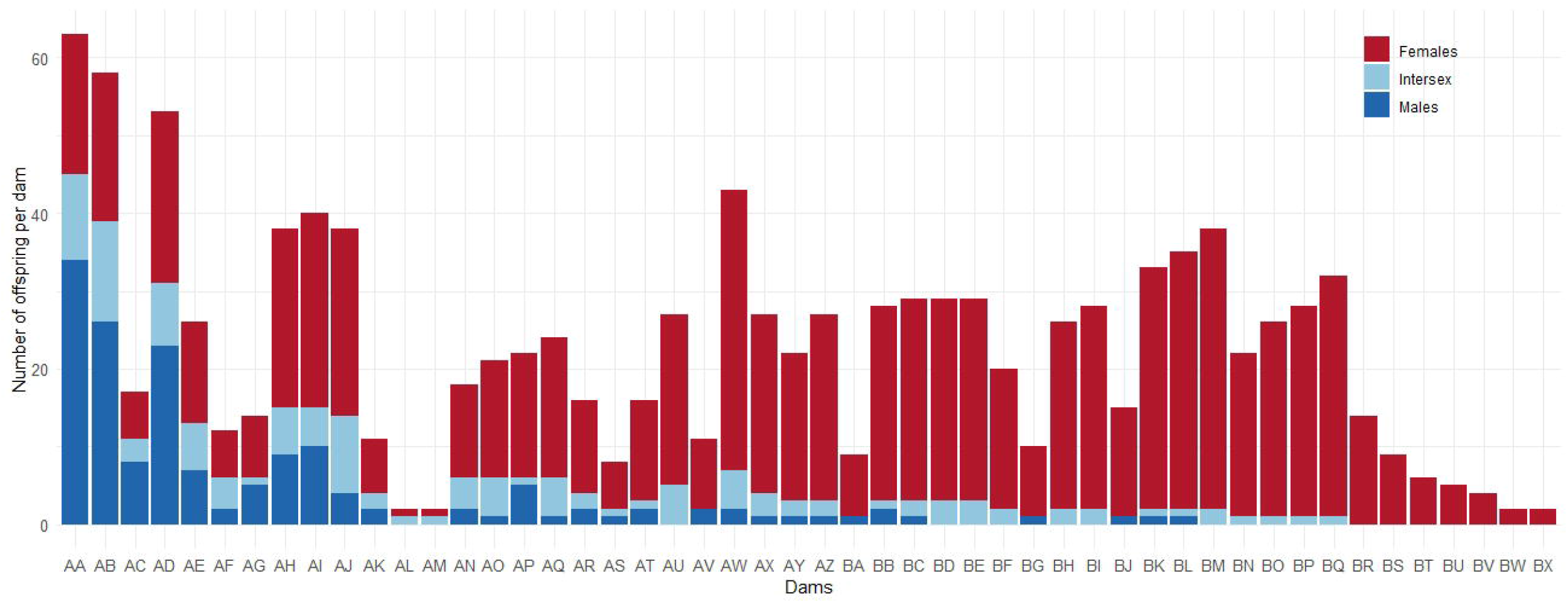
Total number of genotyped offspring (females, males and intersex) for each of the 50 dams (AA to BX) used in the mating scheme. The 22 dams with more than 10 genotyped offspring and extreme masculinisation rate selected for in-depth analysis of their genotypes at QTL regions are underlined.

## Results

### Phenotyping, genotyping and pedigree

In this experiment we phenotyped 20,210 XX-rainbow trout from the French trout farm « *Les Fils de Charles Murgat* ». Among them 98.1% were females (Table 2) and only 1.4% of fish were partially (intersex), or completely masculinised. From the 294 masculinised individuals, 45.0% were intersex with both female and male gonadal tissues. Among these intersex individuals, 60.6% had a right gonad completely masculinised or more masculinised than the left gonad. A significant (p = 3.201e-12) higher maleness (males + intersex) was observed in the group that was reared at 12° (2.0%) than in the group reared at 18° (0.9%) (see Table 2).

**Table 2.** Phenotyping records for the XX-rainbow trout produced at “*Les fils de Charles Murgat*” farm. mpf: months post-fertilisation

| **Temperature treatment (°C) and age at phenotyping** | **12°**<br>**10 mpf** | **12°**<br>**15 mpf** | **Total**<br>**12°** | **18°**<br>**10 mpf** | **18°**<br>**15 mpf** | **Total**<br>**18°** | **Total** |
| --- | --- | --- | --- | --- | --- | --- | --- |
| **Total number of phenotyped fish** | **6,560** | **3,456** | **10,016** | **6,681** | **3,513** | **10,194** | **20,210** |
| **Total number of genotyped fish** | **415** | **353** | **768** | **41** | **343** | **384** | **1152** |
| Phenotyped (and genotyped) females | 6,435<br>(305) | 3,336<br>(258) | **9,771**<br>**(563)** | 6,616<br>(0) | 3,445<br>(295) | **10,061**<br>**(295)** | **19,832**<br>**(858)** |
| Phenotyped and genotyped intersex | 45 | 33 | **78** | 24 | 30 | **54** | **132** |
| Phenotyped and genotyped males | 65 | 62 | **127** | 17 | 18 | **35** | **162** |
| Phenotyped undetermined individuals | 15 | 25 | **40** | 24 | 20 | **44** | **84** |
| Proportion of masculinised individuals (intersex + males) | 1.68% | 2.75% | **2.04%** | 0.61% | 1.37% | **0.87%** | **1.45%** |

From the 1,152 genotyped XX-rainbow trout, 1,139 fish genotyped for 30,811 SNPs were retained after quality controls. Pedigree was recovered for all genotyped offspring except four, the 50 dams had in average 22.7 genotyped offspring (min = 2, max = 63). The average percentage of masculinised fish in the genotyped offspring sample was 20.7%, ranging from 0% for eight females to 71.4% for the AA-dam (see Figure 1 and Supplementary Table S2). The 11 dams (labelled from dam AA to dam AK) that had the highest proportion of masculinised offspring in the genotyped sample bred 370 fish accounting for 68% of masculinised genotyped offspring. On contrary, the 11 dams (labelled from dam BH to dam BR) with a low masculinised rate among their genotyped offspring sample accounted for only 5% of masculinised genotyped offspring. Among those 22 dams, we detected two couples of related (IBD coefficients of 0.36) dams (AA-BK and AI-BR) with one “sister” having a highly masculinised offspring (71.4% for AA or 37.5% for AI) and the other “sister” that produce a low number of masculinised offspring (6.1% for BK or 0% for BR).

### Estimation of genetic parameters for spontaneous maleness in XX-rainbow trout

The variance components and heritability of the sex phenotype estimated under different genomic models using either the 30,811 SNPs on the chip or the 8,784,147 SNPs on WGS are presented in Table 3. The estimated heritability of maleness was high, ranging from 0.48 with the GCTA-chip analysis to 0.62 with the GCTA-seq analysis. The estimated genetic variance (σ^2^_A_) was consistent across all models, with the lower value obtained for GCTA-chip (0.21), intermediate values obtained with BayesCπ models (0.24 and 0.25 for BCπ-chip and BCπ-seq, respectively) and the highest genetic variance estimate (0.26) was obtained with the GCTA-seq analysis. With the increasing number of SNPs (from 31K to WGS) in the model or with the addition of a polygenic effect under a Bayesian approach, the estimate of genetic variance was increased, showing that the 31K SNPs genotyped were insufficient to capture all the genetic variation. Therefore, the real heritability of maleness is expected to be about 0.6 in rainbow trout.

**Table 3.**
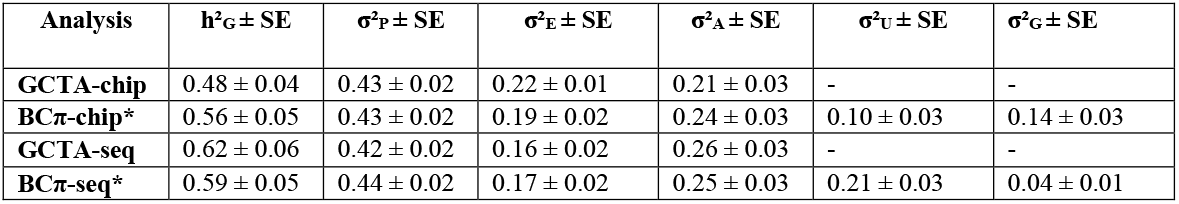
Genetic and genomic parameters estimates for spontaneous maleness under the different statistical models. h^2^_G_ genomic heritability, calculated as σ^2^_A_ / (σ^2^_A_ + σ^2^_E_). σ^2^_A_: Total genetic variance, σ^2^_U_: polygenic variance, σ^2^_G_: genetic variance explained by SNPs, σ^2^_E_: residual variance, σ^2^_P_: phenotypic variance (= σ^2^_A_ + σ^2^_E_). *value of one string among the two or four used for GWAS, the string with the closest final π to the 1% or 0.02% was chosen.

| Analysis | $h^2_G \pm SE$ | $\sigma^2_P \pm SE$ | $\sigma^2_E \pm SE$ | $\sigma^2_A \pm SE$ | $\sigma^2_U \pm SE$ | $\sigma^2_G \pm SE$ |
| --- | --- | --- | --- | --- | --- | --- |
| GCTA-chip | $0.48 \pm 0.04$ | $0.43 \pm 0.02$ | $0.22 \pm 0.01$ | $0.21 \pm 0.03$ | - | - |
| BCπ-chip* | $0.56 \pm 0.05$ | $0.43 \pm 0.02$ | $0.19 \pm 0.02$ | $0.24 \pm 0.03$ | $0.10 \pm 0.03$ | $0.14 \pm 0.03$ |
| GCTA-seq | $0.62 \pm 0.06$ | $0.42 \pm 0.02$ | $0.16 \pm 0.02$ | $0.26 \pm 0.03$ | - | - |
| BCπ-seq* | $0.59 \pm 0.05$ | $0.44 \pm 0.02$ | $0.17 \pm 0.02$ | $0.25 \pm 0.03$ | $0.21 \pm 0.03$ | $0.04 \pm 0.01$ |

Based on the BCπ-chip model, we estimated that the 31K SNPs spanning all the genome accounted for only 58% of the total genetic variance of sex (Table 3). Using the BCπ-seq model we estimated that the sequence segment spanning the 4 Mb located between 62 and 66 Mb on Omy1 accounted for 16% of the total genetic variance of sex (Table 3).

### Genome Wide Association Studies of maleness in XX-female rainbow trout

#### GCTA-chip and BCπ-chip approaches at the 31K genotyping level

Based on the 31K SNPs, we detected four QTLs associated with maleness on three different chromosomes (Omy1, 12 and 20) with the GCTA and the BayesCπ analyses (see Table 4). With the GCTA-chip approach, we detected two QTLs on Omy1, with the first QTL being suggestive only, at 1% at the chromosome-wide level, and explaining less than 0.2% of the total genetic variance. The second QTL was significant, at 1% at the genome-wide level, under GCTA-chip analysis (-log_10_ (P-value) = 11.1) and had a very strong evidence under a BCπ-chip analysis (logBF = 11.3) (Table 4). Even if the two peak SNPs of this QTL differed across the two GWAS approaches with the GCTA peak SNP being located 212 kb before the BCπ peak SNP, they were in close vicinity with only three markers in-between the two peak SNPs. This second QTL explained 3.9% of the total genetic variance. We did not distinguish the two QTLs with the BCπ-chip approach, as the credibility interval estimated with BCπ-chip was 1.7 Mb and contained the two peak SNPs detected with GCTA-chip.

**Table 4.** Summary of QTLs associated with spontaneous maleness in XX-rainbow trout: characteristics for QTLs detected with GCTA-chip and BCπ-chip methods. The QTL start and end positions correspond to the confidence and credibility intervals for GCTA-chip and BCπ-chip, respectively. For BCπ-chip, the logBF is calculated as twice the natural logarithm of Bayes Factor. The % of genetic variance, σ^2^_A_, explained by the QTL was calculated as the sum of the variance by BCπ-chip model of all SNP in the QTL credibility interval divided by the corresponding σ^2^_A_.

| Chromosome | QTL start (Mb) | QTL end (Mb) | Peak SNP name | Peak SNP position (Mb) | Peak $-\log_{10}(P)$ or logBF | % $\sigma^2_A$ explained by QTL | Methods |
| --- | --- | --- | --- | --- | --- | --- | --- |
| 1 | 62.93206 | 63.73809 | Affx-88953453 | 63.53900 | 7.9 | - | GCTA-chip |
| 1 | 64.42011 | 64.63252 | Affx-88916383 | 64.42011 | 11.1 | - | GCTA-chip |
| 1 | 63.45244 | 65.16399 | Affx-88950822 | 64.63252 | 11.3 | 3.86% | BCπ-chip |
| 12 | 6.10355 | 6.79990 | Affx-88940013 | 6.79990 | 8.1 | 0.62% | BCπ-chip |
| 20 | 31.35239 | 31.35239 | Affx-88916019 | 31.35239 | 8.2 | 0.40% | BCπ-chip |

The two other QTLs located on Omy12 and Omy20 were detected using the BCπ-chip approach only (Table 4). The QTL located on Omy12 explained 0.6% of the total genetic variance. The QTL on Omy20, defined by a single SNP located at 31.352 Mb, explained 0.4% of the total genetic variance. This last QTL was suggestive at 5% at the chromosome-wide level under GCTA-chip analysis (-log_10_(P-value) = 4.2).

#### GCTA-seq and BCπ-seq approaches at the whole genome sequence level

Using the WGS data and GCTA-seq approach, we only detected significant QTLs on Omy1, in a delimited region in-between 62 and 66 Mb. In total six peak SNPs corresponding to six putative QTLs (see Supplementary Table S1 and Supplementary Figure S2) were detected. Therefore, the BCπ-seq approach was performed considering only the 21K SNP spanning this 4 Mb-window on Omy1 in addition to a genome-wide polygenic component. Using this approach we detected only two QTLs (see Table 5 and Supplementary Figure S3). The first putative QTL detected by the GCTA-seq analysis was not detected using the BCπ-seq approach. Therefore, it will not be discussed any further as it is likely that the association is only due to linkage disequilibrium with the SNPs included in the second QTL region. The second putative QTL detected between 63.229 and 63.774 Mb with the GCTA-seq approach was also detected with the BCπ-seq approach in a reduced credibility interval (between 63.459 and 63.556 Mb, see Table 5); it was hereafter considered as the first true QTL on Omy1.

**Table 5.** Summary characteristics for the two QTLs associated with maleness in XX-rainbow trout detected on the whole genome sequence using GCTA-seq and BCπ-seq methods. For GCTA-seq, the start and stop position of the first QTL correspond to a CI estimated with a drop-off approach and the start and stop positions of the second QTL correspond to the credibility interval estimated as the union of confidence intervals of four QTLs. The BCπ-seq1 and BCπ-seq2 correspond to the same analysis run with two different seeds for MCMC initialization (random number generator). Significance of peak SNP are calculated as the −log_10_(p-value) for GCTA-seq, and as logBF estimated as twice the natural logarithm of Bayes Factor for BCπ-seq. % σ^2^_A_: proportion of total genetic variance explained by the credibility interval of the QTL calculated using BCπ-seq approaches.

| Chromosome | QTL start (Mb) | QTL end (Mb) | QTL size (kb) | Peak SNP name | Peak SNP position (Mb) | Peak SNP - $\log_{10}(P)$ or logBF | % $\sigma^2_A$ explained by QTL | Methods |
| --- | --- | --- | --- | --- | --- | --- | --- | --- |
| 1 | 63.229379 | 63.774478 | 545.1 | Omy1-63493395 | 63.493395 | 10.61 | - | GCTA-seq |
| 1 | 64.360291 | 64.707100 | 341.6 | Omy1-64597800 | 64.597800 | 13.74 | - | GCTA-seq |
| 1 | 63.459553 | 63.555809 | 96.27 | Omy1-63520900 | 63.542090 | 12.97 | 0.56% | BCπ-seq1 |
| 1 | 64.266321 | 64.649580 | 383.26 | Omy1-64607018 | 64.607018 | 14.67 | 13.55% | BCπ-seq1 |
| 1 | 63.459545 | 63.727071 | 267.53 | Omy1-63520900 | 63.542090 | 11.55 | 0.51% | BCπ-seq2 |
| 1 | 64.087947 | 64.632497 | 544.55 | Omy1-64610233 | 64.610233 | 17.32 | 14.56% | BCπ-seq2 |

The four remaining putative QTLs from the GCTA-seq analysis were very close to each other, with less than 250 kb between two successive peak SNPs, and the limits of their confidence intervals (CIs) approximated by a drop-off approach were all distant from less than 3 kb. Therefore these four CIs were fused into a single credibility interval of 342 kb. This second QTL was also detected with the BCπ-seq analysis (Table 5). While the peak SNP of the first QTL was the same under the two BCπ-seq analyses, this was not the case for this second QTL. However, the two peak SNPs were only separated by 3.2 kb and the credibility intervals were strongly overlapping (Table 5).

Based on the average of the two BCπ-seq analyses, among the 16% of genetic variance explained by all the 21K SNPs included in the BCπ-seq model, the first QTL explained about 0.5% and the second QTL explained about 14% of the genetic variance.

At the end of the second QTL we found a haplotype block of 15 consecutive SNPs (spanning 745 kb from 64.632011 to 64.632756 Mb) that have a significant effect on masculinisation (all the 15 SNPs have −log_10_(P-value) between 12.2 and 13.6 and among them five SNPs with logBF > 9). For this 15 SNP-haplotype, masculinised fish were overrepresented among the homozygous genotypes for the alternative allele (different from the reference genome sequence). Interestingly, for the same block of 15 SNPs, eight of the 11 dams with almost no masculinised offspring carried two copies of the reference genome haplotype (the three remaining dams being heterozygous at these 15 SNPs). Among the 11 dams with a high rate of masculinised offspring, only three were homozygous for the reference haplotype, six were heterozygous and the last two (labelled AA and AJ in Figure 1) were homozygous for the alternative haplotype. In the offspring, the homozygous genotype for the alternative haplotype was overrepresented in masculinised fish (51.5% of masculinised fish) and underrepresented in females (8.7% of females). On the contrary, the reference haplotype was observed either at the heterozygous or homozygous state for respectively 50.6% and 40.2% of the female progeny.

### Positional candidate genes and SNP annotation

In total five genes were positioned within the first QTL region spanning from 63.459 to 63.556 Mb (see Supplementary Table S3). The peak SNP from GCTA-seq (at 63.493 Mb) was located within the *pygb* gene (glycogen phosphorylase, brain form, from 63.487 to 63.508 Mb), and the peak SNP from both BCπ-seq runs (6.542 Mb) was located within the *ninl* gene (ninein-like protein, from 63.542 to 63.594 Mb). Among the 669 annotated SNPs spanning the QTL region, only 23 SNPs were indicated with either a low (16 SNPs) or a moderate (7 SNPs) potential effect on gene expressions when annotated with the SNPEff software (See Figure 2 and Supplementary Table S4). The only SNP that was both significant (-log_10_(P-value) = 9.6) and annotated with a potential moderate effect was located at 63 543 061 bp within the *ninl* gene and annotated has a missense variant (Figure 2). This SNP was very close to the peak SNP detected by the two BCπ-seq analyses (Table 5).

**Figure 2.**
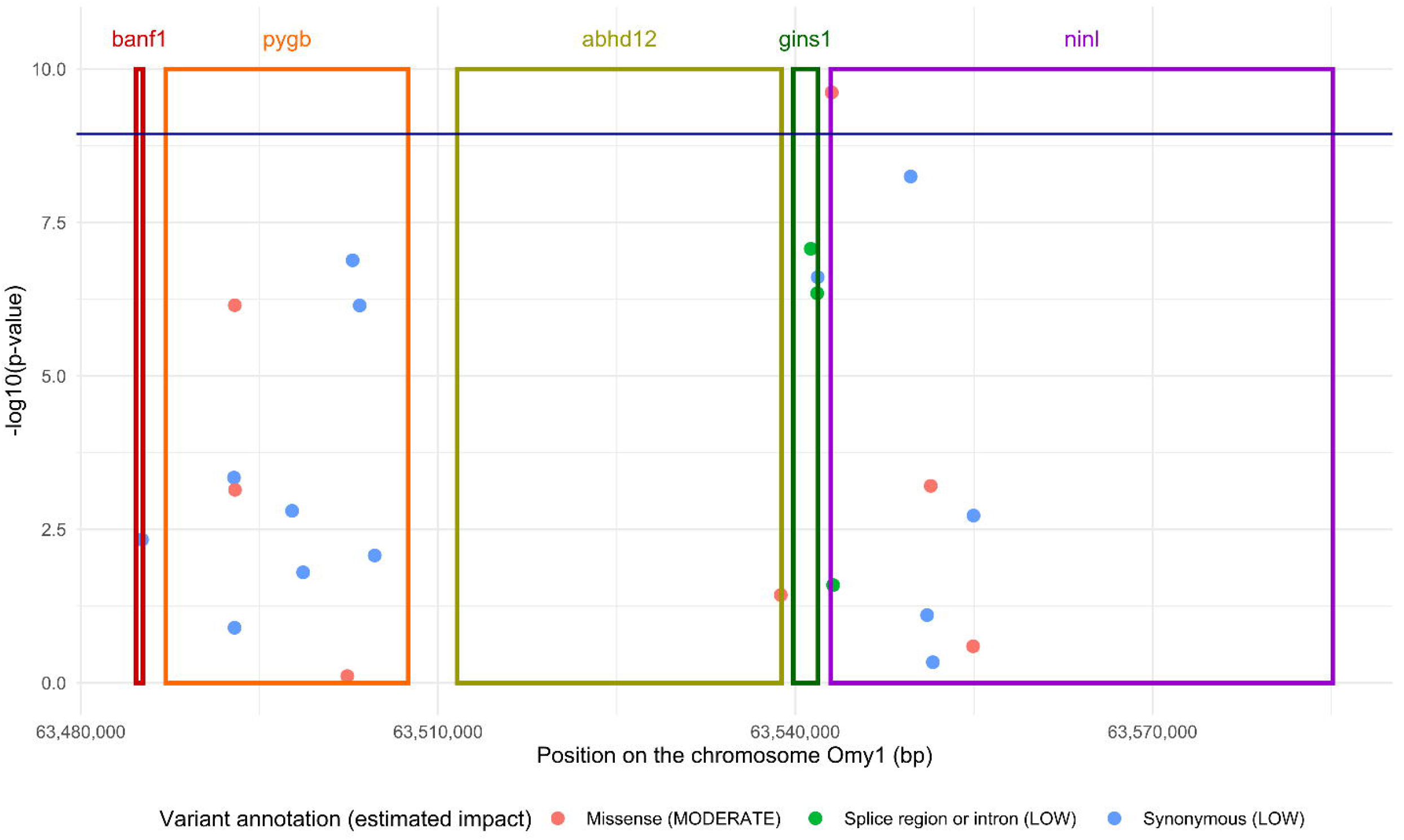
Annotated SNPs located within the first QTL region from 63.549 to 63.556 Mb on the chromosome 1. The significance of SNP (dots) effect is given by the −log_10_(p-value) estimated with the GCTA-seq approach. The dark blue is the 1% at the genome wide level threshold. Putative impact of SNP on genes was estimated by SNPEff, only SNP with MODERATE or LOW effects on genes are represented. Genes located within this QTL are represented by rectangle: *banf1* (barrier-to-autointegration factor-like), *pygb* (glycogen phosphorylase, brain form), *abhd12* (alpha/beta-Hydrolase domain containing 12), *gins1* (DNA replication complex GINS protein PSF1), *ninl* (ninein-like protein).

Within the second QTL region spanning from 64.360 to 64.633 Mb, 14 genes were annotated. All the 3 peak SNPs (table 5) detected by the different WGS analyses were located in the intergenic region between the *hells* gene (helicase, lymphoid specific) and an uncharacterized protein (LOC110527930, see Supplementary Table S3). In total, 1,498 SNPs were annotated in the QTL region, and only 44 SNPs had either a moderate (15 SNPs) or low (29 SNPs) potential effect on gene expression (Figure 3 and Supplementary table E). Among those 44 SNPs, 12 SNPs with a potential effect on 3 genes i.e., *cyp17a1* (Cytochrome P450 Family 17 Subfamily A Member 1), *hells*, and LOC110527930 were significant (-log_10_(P-value) >10) based on GCTA-seq analysis (Figure 3). One significant SNP (-log_10_(P-value) = 10.2) was annotated as a missense variant within the *cyp17a1* gene (Figure 3). Two significant SNPs (-log_10_(P-value) = 12.8 and 11.7) were annotated for a low potential effect on the *hells* gene as they corresponded respectively to a synonymous mutation and a variant within a splice region or an intron of the gene (see Supplementary Table S5). Finally, nine SNPs from the haplotype block of 15 SNPs previously identified, were annotated with either low or moderate potential effects on the LOC110527930 uncharacterized protein (see Supplementary Table S5). All the seven SNPs annotated for moderate effects were missense variants. Among them, two SNPs, located at 64 632 546 bp and at 64 632 583 bp (Figure 3), seemed to be good candidates for the causative mutation as they were both significant in the GCTA-seq and one BCπ-seq analysis (-log_10_(P-value) = 13.6 and logBF > 11).

**Figure 3.**
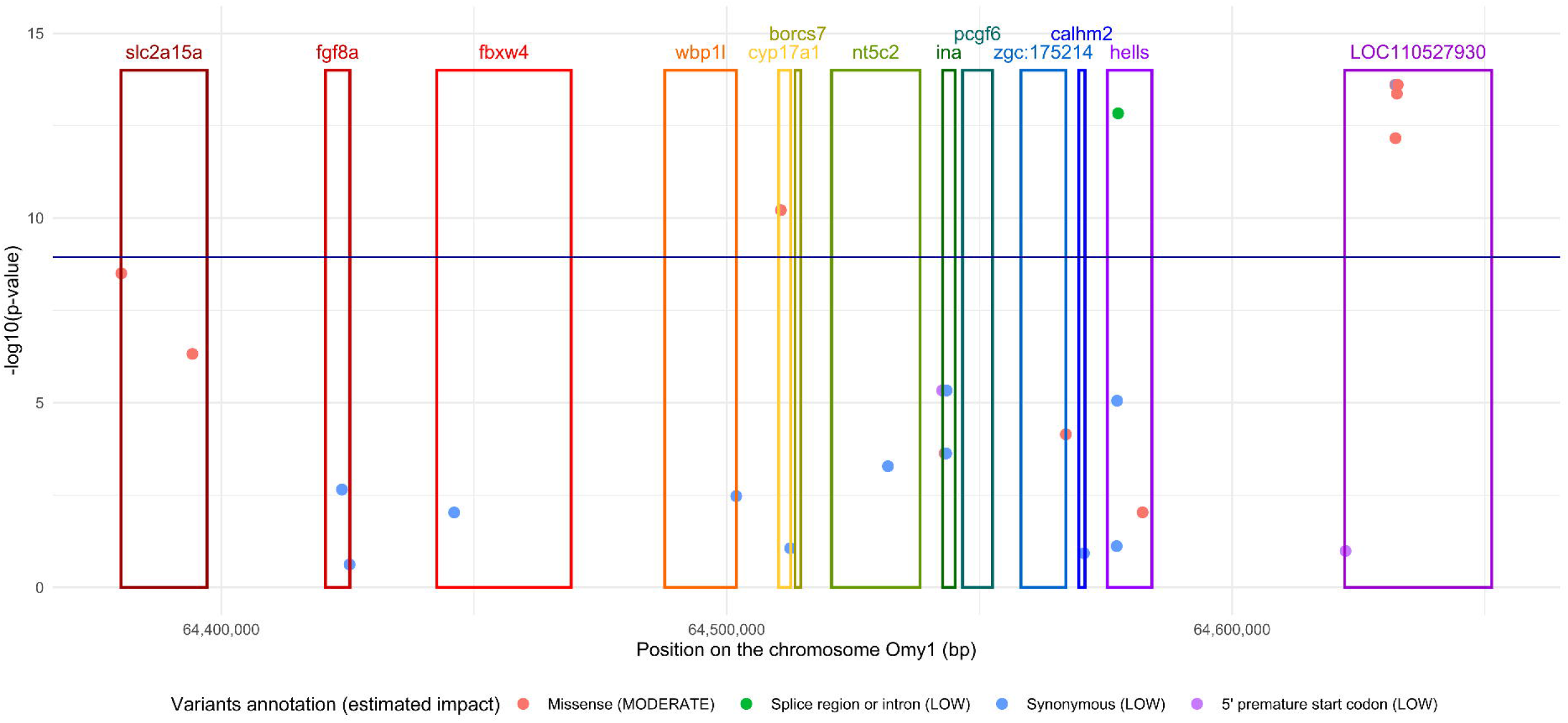
Annotated SNPs located within the second QTL region, from 64.360 to 64.633 Mb on the chromosome 1. The significance of SNPs (dots) is given by the −log_10_(p-value) estimated with the GCTA-seq approach. The dark blue is the 1% at the genome wide level threshold. Putative impact of SNP on genes was estimated by SNPEff. Only SNP with MODERATE or LOW effects on genes are represented. Genes located within this QTL are represented by rectangle: *slc2a15a* (solute carrier family 2, facilitated glucose transporter member 9-like), *fgf8* (fibroblast growth factor 8), *fbxw4* (F-box/WD repeat-containing protein 4), *wbp1l* (WW domain binding protein 1-like), *cyp17a1* (Cytochrome P450 Family 17 Subfamily A Member 1), *borcs7* (BLOC-1-reltaed complex subunit 7), *nt5c2* (cytosolic purine 5’-nucleotidase), *ina* (alpha-internexin), *pcgf6* (polycomb group RING finger protein 6), *zgc:175214* (RING finger protein 122-like), *calhm2* (calcium homeostasis modulator protein 2), *hells* (lymphocyte-specific helicase-like), LOC110527930 (uncharacterized protein).

## Discussion

To the best of our knowledge, this study is the first one to study spontaneous masculinisation in XX rainbow trout through a GWAS. The analysis was carried out in a rainbow trout commercial line reared either under standard temperatures (12-14.5°C) or after a high temperature treatment during early stages.

The overall masculinisation rate was 1.45%, which is in range of what is reported by several French farmers (unpublished data). Unexpectedly, the masculinisation rate was significantly higher for the group reared at the 12°C water temperature than for the group exposed to the high temperature treatment at 18°C. This result is in contradiction with the observation from Valdivia *et al*.^15^ (2014) who observed a masculinizing effect of early hot temperature treatment (18°C vs 12°C). However, in their experiment, the temperature treatment was applied a little bit earlier (at 32 dpf, instead of 35 dpf), so the temperature we applied might have not completely overlap the critical thermosensitive period of sex differenciation^35,36^ inducing a repression instead of an induction of masculinisation. This hypothesis should deserve further work to be confirmed. Different genetic backgrounds could also contribute to the observed differences in response to thermal treatment. In Validivia *et al.^15^* (2014), the tested families belonged to the INRA XX-*mal* lineage, an experimental line different from the commercial population used in this study. The hypothesis is further supported by the fact that distinct maleness QTLs were detected in INRA XX-*mal* families^13^ and in this study (see below), suggesting that masculinizing factors may differ according to populations, resulting in different susceptibility to temperature. Indeed, different sex ratios have been reported in rainbow trout populations with various genetic backgrounds in response to temperature treatments^14,37^. In other fish species such as Atlantic silverside^38^ or Nile tilapia^39^, sex differentiation is also known to be dependent of both the water temperature and the genetic background of the population.

The observed level of masculinisation was variable among masculinised fish with about 40 and 61% of intersex fish in the 12°C and 18°C groups, respectively. Within those intersex individuals, whatever the rearing temperature of fry, the right gonad was very often more masculinised than the left gonad. This particularity was first described by Quillet *et al*. (2004)^40^ and confirmed by Valdivia *et al*. (2013, 2014)^12,15^, in families originating from the INRA XX-*mal* carrying line. Under normal rearing condition, the left-right asymmetry (LR) is almost undetectable as intersex fish remain rare and poorly described. However, the observation of such an asymmetry in another rainbow trout population suggests that this a general feature in rainbow trout and that gonads in this species develop in an asymmetrical LR manner, as it has been reported in mammals, birds, amphibians and reptiles^41^ and in some fish^42–44^.

In this study, we reported high genomic heritability estimates for spontaneous maleness (from 0.48 up to 0.62) with the QTLs detected explaining only a small proportion (up to 14%) of the estimated genetic variance. Using the BCπ-chip model, we estimated that 58% of the genetic variance was explained by the pan-genomic 31K SNPs. The remaining 42% of genetic variance was probably not captured by the SNPs due to their heterogeneous density distribution across the genome and also due the inaccurate estimates (regressed towards 0) of their effects with only 1,139 phenotyped and genotyped individuals.

Concerning the identification of the genetic determinants of the maleness, we detected four QTLs on three different chromosomes, two QTLs being on Omy1, one on Omy12 and the last one on Omy20. Both QTLs on Omy12 and Omy20 were only detected in the 31K SNPs analyses and were not confirmed on WGS analyses. The QTL on Omy20 might be the result of a spurious association as it was defined by a single SNP in both BCπ-chip and GCTA-chip analysis. It is interesting to note that in their study in gynogenetic families from INRA XX-*mal* line, Guyomard *et al.^13^* (2014) detected four QTLs associated with spontaneous maleness on four different chromosomes and that one QTL was located on Omy20 (linkage group RT17) too. However, it had a wide CI, covering the entire chromosome, so it is not possible to say if the QTL we detected in the present study on Omy20 is the same and is shared between the two genetic backgrounds. As we did not detect QTL on the three other chromosomes they reported, maleness in rainbow trout with different genetic backgrounds is likely to be controlled by a variety of underlying genetic mechanisms.

The two main QTLs associated with masculinisation in our population were both detected on Omy1, which has never been reported as associated with neither sex determinism nor maleness in rainbow trout. Using WGS information, we were able to reduce the CI of the first QTL from 806 kb (GCTA-chip, Table 4) to 545 kb based on GCTA-seq analysis and even 96 kb for the credibility interval estimated with one of the BCπ-seq runs (Table 5). While highly significant, this QTL explained only 0.5% of the total genetic variance. This low proportion of variance explained may be the result of a strong linkage disequilibrium of those SNPs with some SNPs of the second QTL, which was mapped at a close distance (< 1.2 Mb between the peak SNPs of these two QTLs), making this first QTL an artefact. Within this first QTL region, no gene seems to have a functional role in gonad development; however, this QTL may contain regulating factors involved in the control of genes contained in the second QTL.

For the second QTL, the credibility interval ranged from 268 kb to 545 kb depending on the different WGS analyses (Table 5 and Supplementary Table S1). Based on the BCπ-seq analysis, we estimated that this second QTL explain about 14% of the total genetic variance while it explained less than 4% of the total genetic variance under the BCπ-chip analysis. It is difficult to know whether this proportion of variance was either overestimated under the BCπ-seq model or underestimated under the BCπ-chip model because we ignored the covariance between close SNPs estimates when deriving this proportion. Nevertheless, this QTL is explaining a sufficient part of the genetic variance to deserve specific attention in breeding programs.

Within this main QTL region, three genes may be potential candidates for explaining the spontaneous masculinisation. First, the *fgf8a* gene might be involved in the left-right (LR) asymmetrical gonad development observed in the intersex fish. This *fgf8* gene is part of the fibroblast growth factor family which plays a major role in vertebrate development and has been reported to be fundamental for proper LR asymmetric development of multiples organs (brain, heart and gut) in zebrafish^45^. The second and biologically more convincing candidate gene is *cyp17a1* as it is a major enzyme of the steroidogenic pathway and is involved in the synthesis of biologically active gonadal androgens and oestrogens. Indeed, *cyp17a1* has been identified as potentially involved in spontaneous maleness in common carp (*Cyprinus carpio*)^46^ and was recently characterized as being involved in the gonad differentiation in zebrafish with complete masculinisation phenotype in *cyp17a1* deficient zebrafish^47^. However, in our analysis, the only SNP with a putative missense effect for this gene was significant in the GCTA-seq analysis only (-log_10_(p-value) = 10.2) and had very low logBF values in both BCπ-seq approach (0 or 2.4). However, even if the *cyp17a1* gene was not within the most significant region of the QTL (in-between the *hells* gene and LOC110527930) in terms of significance/evidence tests, we could not exclude its implication potentially through long distance regulation of its expression. Based on the GWAS as well as the SNP annotation, the uncharacterized LOC110527930 protein was the most relevant candidate found in our analysis with a haplotype block of 15 consecutive SNPs (from Omy1-64632011 to Omy1-64632756) that were all significant (-log_10_(P-value) > 12) and presented alternative genotypes for 4 dams with opposite proportions of masculinised progeny (2 dams with highly masculinised genotyped offspring and 2 dams with low masculinised genotyped offspring). In addition, seven of those SNPs were annotated with a putative missense effect on the LOC110527930 protein. However, there is no published evidence that this uncharacterized protein could play a role in gonadal sex differentiation and more work is still needed to characterise this protein and its potential effect on gonadal differentiation.

The high heritability (up to 0.62) of spontaneous maleness estimated in this study opens up opportunities to manage maleness in all-female trout populations. In particular, in case the two highly significant QTLs detected on Omy1 in the present population were confirmed to play a role in spontaneous maleness in numerous other rainbow trout populations with diverse genetic origins, the identified SNPs could then be used in a cost-efficient genotyping test to identify female broodstock with a higher propensity to transmit male or intersex phenotype in their progeny. Breeders willing to limit the occurrence of undesirable masculinised individuals in their commercial all-female stocks, could target those SNPs to discard broodstock that could sire a masculinised progeny. We tested various combinations of SNPs from the two QTLs detected on Omy1 in order to maximise the efficiency of detection of the masculinised fish while keeping low the number of false positives (*i.e*. females) to develop a test with good specificity in our studied population. Before applying the same test in other populations, the effect of those SNPs should be confirmed in a large set of populations with diverse genetic backgrounds. Our preliminary results regarding maleness QTLs in either the XX-*mal* families from the INRA experimental population and the French commercial population used in this study suggest that this might not be the case. Within the main QTL (from 64.360 to 64.707 Mb), all significant SNPs would give similar results, any SNP homozygous for the alternative allele would allow the detection of 44.7 to 51.5% of masculinised fish in our sample, whereas only 6.8 to 13.7% of the homozygous fish for the alternative allele would be females. In particular, using the alternative allele of any SNP from the haplotype block of 15 SNPs described previously (between 64,632,011 and 64,632,756 bp) would allow detecting 51.5% of masculinised fish in our sample. Among those 15 SNPs, the SNP Affx-88950822 is present on the commercial 57K chip and could be of immediate use, provided its effect is found in other trout populations. Every combination of two or more SNPs within this main QTL would results in a smaller proportion of false positive (less females discarded) but also in a slightly smaller proportion of males identified. Still in our samples, pairing this Affx-88950822 SNP (or any other SNP from the associated 15-SNP haploblock) with SNPs from the first QTL on Omy1 (from 63,459 to 63,556 Mb) would slightly increase the test sensitivity (identifying more masculinised fish) without eliminating too many true females.

From a breeder perspective, additional research is needed to improve knowledge about genetic basis and environmental factors determining spontaneous maleness in all-female stocks before any application in the industry. In addition to the confirmation of the existence of the same QTLs in various rainbow trout populations, the expected genetic and phenotypic responses under different thermal regimes and either pedigree-based or marker-assisted selection should be quantified to assess the efficiency of the strategy to limit the rate of spontaneous males in the production of all-female stocks. Further investigation is also needed with regard to the production of sex-reversed male breeders based on the use of spontaneous sexual inversion to prevent the use of hormones. Indeed, the high heritability of maleness suggests that the use of spontaneously masculinised individuals as progenitors of all-female populations would increase the frequency of undesirable masculinized progeny. Combining masculinising genetic factors together with an environmental (temperature) control of gonad masculinization according to the destination of the fish (broodstock vs all-female production stock) and/or the rearing environment might offer a solution to manage the trade-off. However, because of the overall low masculinization rates recorded in this study, we were not able to detect any potential interaction between rearing temperature and genotype, and we have no suspicion of the existence of such an interaction, as a GWAS with GCTA-chip, carried out with individuals from the 12°C group only, detected the same QTLs as for the overall population (results not shown). In the current state of knowledge on the effect of temperature on sexual differentiation, it is too early to propose an efficient way of controlling the rearing conditions to either enhance or limit spontaneous maleness in rainbow trout. Although the prospect of using hormone-free neomales is very attractive, a potential important drawback may be to come back to the flesh quality and animal welfare issues (flesh loss of lipids, color and firmness due to precocious maturation; mortality due to Sparolegnia fungus) initially encountered in commercial stocks with XY-males.

In conclusion, in this study we detected minor genetic factors involved in maleness in rainbow trout in the absence of the master gene *sdY*. Two QTLs detected on Omy1 explained up to 15% of the total genetic variance of maleness in the population used in this study and we identified three candidate genes that might be involved in the masculinisation of XX-rainbow trout. Among those three genes, one uncharacterised protein (LOC110527930) was the most relevant candidate and more work would be need to characterise the potential effect of this protein on gonadal differentiation.

## Acknowledgements

The authors are grateful to R. Murgat and the staff of “*Les Fils de Charles Murgat*” farm for their help and care of the fish. The authors thank C. Blay, J. D’Ambrosio and F. Enez (SYSAAF), J. Bobe, E. Cardonna, A. Fostier, L. Goardon, A. Herpin, H. Hu, D. Lallias, M. Vandeputte and M. Wen (INRAE) and F. Ruelle (Ifremer) for their helpful participation to the recording of phenotypes.

The European Maritime and Fisheries Fund and FranceAgrimer co-funded this work (NeoBio project, n° R FEA470016FA1000008).

## Author contributions statement

C.F. performed the data curation, interpretation and formal analysis, prepared all figures and wrote the original draft. F.P was involved in the conceptualisation and supervision of the project, the development of statistical methodology and data interpretation and wrote the original draft. A.B., C.P., P.H., F.K., N.D., C.C., Y.G. and EQ. were involved in animal sampling and phenotyping. F.K., N.D., C.C., M.M, J.L. and O.B. were involved in laboratory works. M.C., M.B, C.H. and F.P. were involved in the data curation, and helped with programming and software utilisation. A.B., C.P., P.H., P.H. and Y.G. were involved in the conceptualisation of the project, in funding acquisition and in the design of the protocol. E.Q. was project coordinator involved in the conceptualisation and coordination of the whole study, funding acquisition, design of the protocol, data interpretation and revision of the original draft. All authors reviewed and approved the manuscript.

## Additional information

### Data availability

The datasets for this manuscript are not publicly available because data belongs partly to a private company. The data can be made available for reproduction of the results from Florence Phocas and Charles Murgat Pisciculture on request via a material transfer agreement.

### Supplementary information

Supplementary Table S1. GCTA-seq QTL definition with their confidence intervals.

Supplementary Table S2. Descriptive statistics of number of offspring according to their phenotypic sex and proportion of masculinised offspring per dams

Supplementary Table S3. Candidate genes. QTLs start and end correspond to a reduced interval determined as the intersection of confidence and credibility intervals estimated using both GCTA-seq and BCπ-seq analysis.

Supplementary Table S4. Annotated SNPs within the first QTL region. Only SNPs with a moderate or low annotated impact are presented.

Supplementary Table S5. Annotated SNPs within the second QTL region. Only SNPs with a moderate or low annotated impact are presented.

**Supplementary Figure S1.**
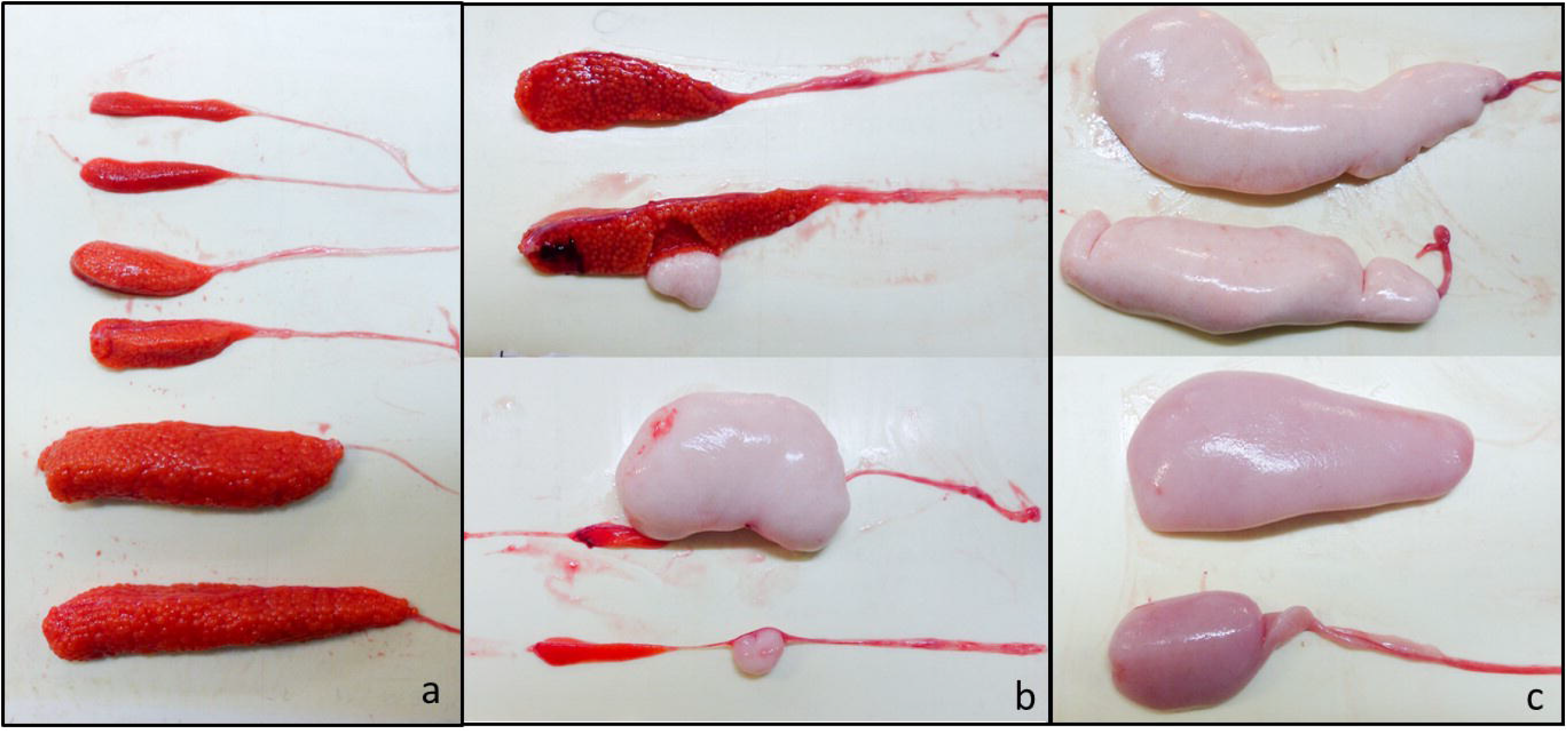
Photos of a) female, b)intersex and c)male gonads from phenotyped fish.

**Supplementary Figure S2.**
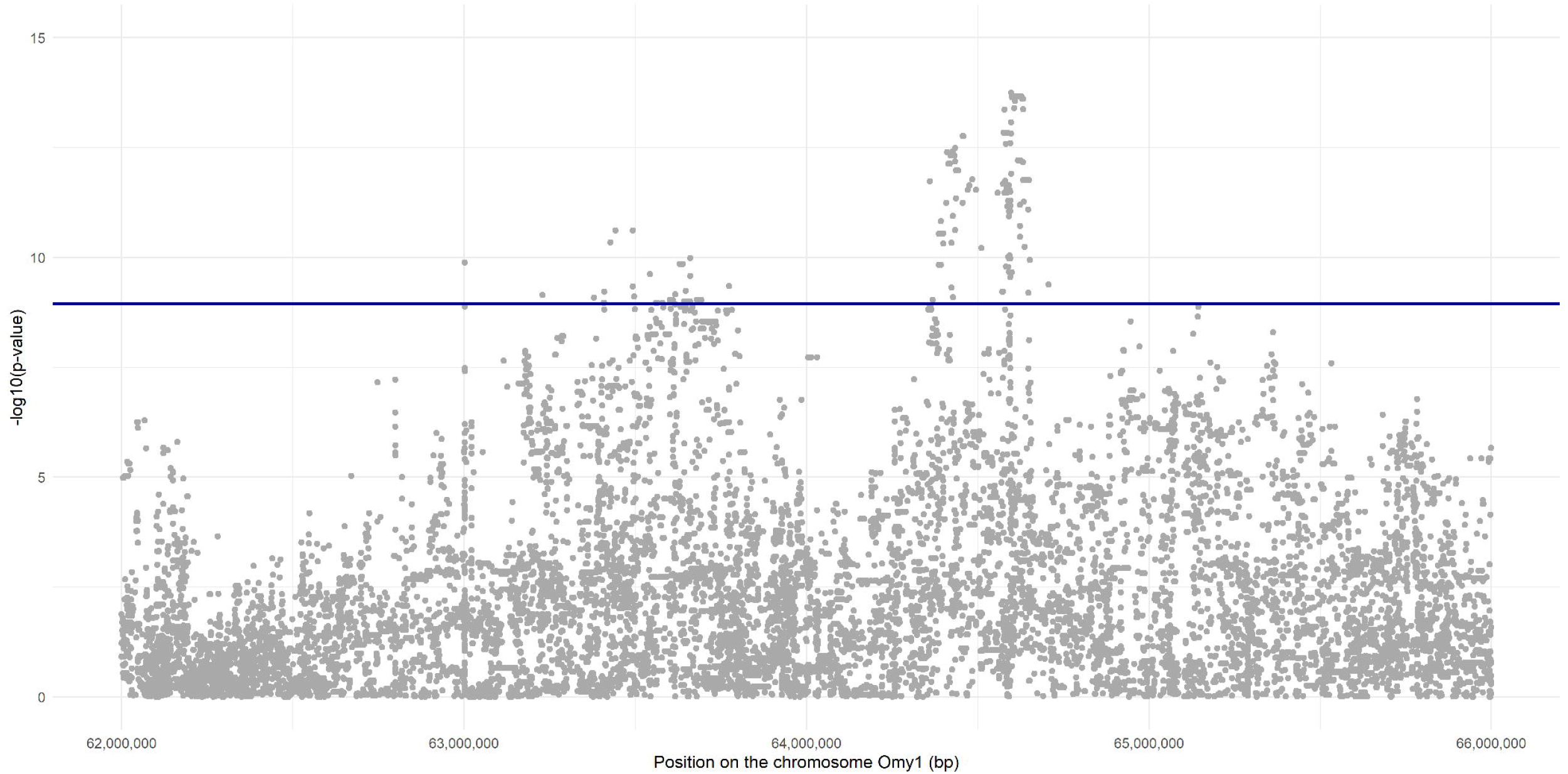
Manhattan plot for the GCTA-seq analysis for SNPs located between 62 and 66 Mb in the chromosome Omy1. The dark blue line is the 1% genome wide threshold estimated with Bonferroni correction.

**Supplementary Figure S3.**
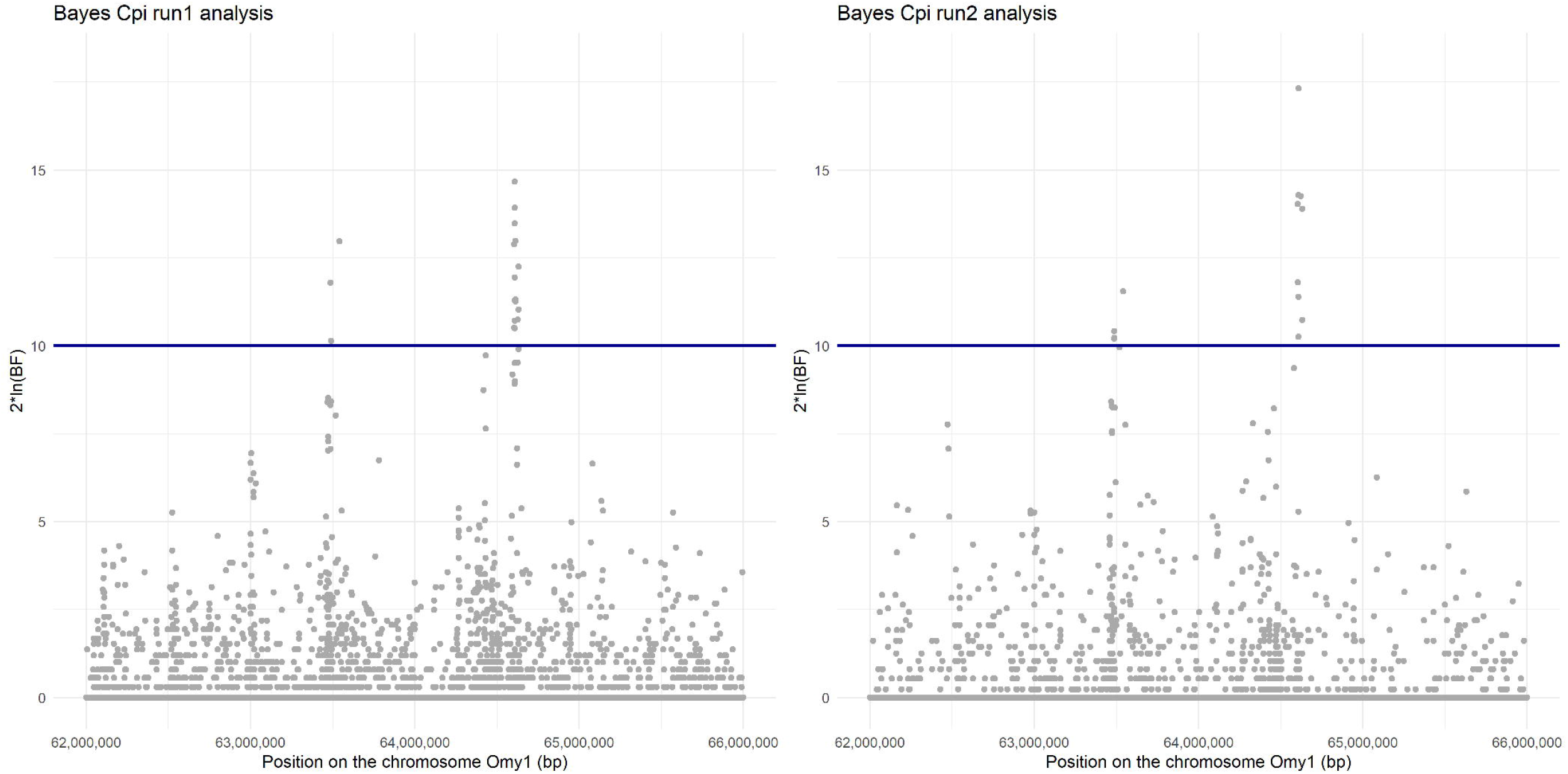
Manhattan plots for the two BCπ-seq analysis performed with two seeds. The dark blue line correspond to the very strong evidence in favour of a QTL threshold. BF = Bayes Factor

## References

1. Devlin, R. H. & Nagahama, Y. Sex determination and sex differentiation in fish: an overview of genetic, physiological, and environmental influences. Aquaculture 208, 191–364 (2002).

2. Martínez, P. et al. Genetic architecture of sex determination in fish: applications to sex ratio control in aquaculture. Front. Genet. 5, (2014).

3. Mank, J. E. & Avise, J. C. Evolutionary diversity and turn-over of sex determination in teleost fishes. Sex. Dev. Genet. Mol. Biol. Evol. Endocrinol. Embryol. Pathol. Sex Determ. Differ. 3, 60–67 (2009).

4. Johnstone, R., Simpson, T. H., Youngson, A. F. & Whitehead, C. Sex reversal in salmonid culture: Part II. The progeny of sex-reversed rainbow trout. Aquaculture 18, 13–19 (1979).

5. Chevassus, B., Devaux, A., Chourrout, D. & Jalabert, B. Production of YY rainbow trout males by self-fertilization of induced hermaphrodites. J. Hered. 79, 89–92 (1988).

6. Chourrout, D. & Quillet, E. Induced gynogenesis in the rainbow trout: Sex and survival of progenies production of all-triploid populations. Theor. Appl. Genet. 63, 201–205 (1982).

7. Yano, A. et al. An Immune-Related Gene Evolved into the Master Sex-Determining Gene in Rainbow Trout, Oncorhynchus mykiss. Curr. Biol. 22, 1423–1428 (2012).

8. Yano, A. et al. The sexually dimorphic on the Y-chromosome gene (sdY) is a conserved male-specific Y-chromosome sequence in many salmonids. Evol. Appl. 6, 486–496 (2012).

9. Bertho, S. et al. Foxl2 and Its Relatives Are Evolutionary Conserved Players in Gonadal Sex Differentiation. Sex. Dev. Genet. Mol. Biol. Evol. Endocrinol. Embryol. Pathol. Sex Determ. Differ. 10, 111–129 (2016).

10. Bertho, S. et al. The unusual rainbow trout sex determination gene hijacked the canonical vertebrate gonadal differentiation pathway. Proc. Natl. Acad. Sci. U. S. A. 115, 12781–12786 (2018).

11. Quillet, E., Aubard, G. & Quéau, I. Mutation in a sex-determining gene in rainbow trout: detection and genetic analysis. J. Hered. 93, 91–99 (2002).

12. Valdivia, K. et al. Sex Differentiation in an All-Female (XX) Rainbow Trout Population with a Genetically Governed Masculinization Phenotype. Sex. Dev. 7, 196–206 (2013).

13. Guyomard, R. et al. RAD-seq mappingof spontaneous masculinization in XX doubled haploid rainbow trout lines. in (2014).

14. Magerhans, A., Müller-Belecke, A. & Hörstgen-Schwark, G. Effect of rearing temperatures post hatching on sex ratios of rainbow trout (Oncorhynchus mykiss) populations. Aquaculture 294, 25–29 (2009).

15. Valdivia, K. et al. High Temperature Increases the Masculinization Rate of the All-Female (XX) Rainbow Trout “Mal” Population. PLOS ONE 9, e113355 (2014).

16. Ospina-Álvarez, N. & Piferrer, F. Temperature-Dependent Sex Determination in Fish Revisited: Prevalence, a Single Sex Ratio Response Pattern, and Possible Effects of Climate Change. PLOS ONE 3, e2837 (2008).

17. Baroiller, J. F., D’Cotta, H. & Saillant, E. Environmental effects on fish sex determination and differentiation. Sex. Dev. Genet. Mol. Biol. Evol. Endocrinol. Embryol. Pathol. Sex Determ. Differ. 3, 118–135 (2009).

18. Jalabert, B., Billard, R., Chevassus, B., Escaffre, A. & Carpentier, M. Preliminary experiments on sex control in trout: production of sterile fishes ans dimultaneous self-fertilizable hermaphrodites. Ann. Biol. Anim. Biochim. Biophys. 15, 19–28 (1975).

19. Palti, Y. et al. The development and characterization of a 57K single nucleotide polymorphism array for rainbow trout. Mol. Ecol. Resour. 15, 662–672 (2015).

20. D’Ambrosio, J. et al. Genome-wide estimates of genetic diversity, inbreeding and effective size of experimental and commercial rainbow trout lines undergoing selective breeding. Genet. Sel. Evol. 51, 26 (2019).

21. Sargolzaei, M., Chesnais, J. P. & Schenkel, F. S. A new approach for efficient genotype imputation using information from relatives. BMC Genomics 15, (2014).

22. Gao, G. A New and Improved Rainbow Trout (Oncorhynchus mykiss) Reference Genome Assembly. in vol. 1 40 (2016).

23. Danecek, P. et al. The variant call format and VCFtools. Bioinformatics 27, 2156–2158 (2011).

24. Chang, C. C. et al. Second-generation PLINK: rising to the challenge of larger and richer datasets. GigaScience 4, 7 (2015).

25. Yang, J., Lee, S. H., Goddard, M. E. & Visscher, P. M. GCTA: A Tool for Genome-wide Complex Trait Analysis. Am. J. Hum. Genet. 88, 76–82 (2011).

26. Yang, J. et al. Common SNPs explain a large proportion of the heritability for human height. Nat. Genet. 42, 565–569 (2010).

27. Habier, D., Fernando, R. L., Kizilkaya, K. & Garrick, D. J. Extension of the bayesian alphabet for genomic selection. BMC Bioinformatics 12, 186 (2011).

28. Boerner, V. & Tier, B. BESSiE: a software for linear model BLUP and Bayesian MCMC analysis of large-scale genomic data. Genet. Sel. Evol. GSE 48, 63 (2016).

29. Li, H. A quick method to calculate QTL confidence interval. J. Genet. 90, 355–360 (2011).

30. Schurink, A., Janss, L. L. & Heuven, H. C. Bayesian Variable Selection to identify QTL affecting a simulated quantitative trait. BMC Proc. 6, S8 (2012).

31. Kass, R. E. & Raftery, A. E. Bayes Factors. J. Am. Stat. Assoc. 90, 773–795 (1995).

32. Vidal, O. et al. Identification of carcass and meat quality quantitative trait loci in a Landrace pig population selected for growth and leanness. J. Anim. Sci. 83, 293–300 (2005).

33. Michenet, A., Barbat, M., Saintilan, R., Venot, E. & Phocas, F. Detection of quantitative trait loci for maternal traits using high-density genotypes of Blonde d’Aquitaine beef cattle. BMC Genet. 17, 88 (2016).

34. Cingolani, P. et al. A program for annotating and predicting the effects of single nucleotide polymorphisms, SnpEff: SNPs in the genome of Drosophila melanogaster strain w ^1118^; iso-2; iso-3. Fly (Austin) 6, 80–92 (2012).

35. Azuma, T. et al. The influence of temperature on sex determination in sockeye salmon Oncorhynchus nerka. Aquaculture 234, 461–473 (2004).

36. Craig, J. K., Foote, C. J. & Wood, C. C. Evidence for temperature-dependent sex determination in sockeye salmon (Oncorhynchus nerka). Can. J. Fish. Aquat. Sci. 53, 141–147 (1996).

37. Magerhans, A. & Hörstgen-Schwark, G. Selection experiments to alter the sex ratio in rainbow trout (Oncorhynchus mykiss) by means of temperature treatment. Aquaculture 306, 63–67 (2010).

38. Conover, D. O. & Kynard, B. E. Environmental sex determination: interaction of temperature and genotype in a fish. Science 213, 577–579 (1981).

39. Baroiller, J. F., D’Cotta, H., Bezault, E., Wessels, S. & Hoerstgen-Schwark, G. Tilapia sex determination: Where temperature and genetics meet. Comp. Biochem. Physiol. A. Mol. Integr. Physiol. 153, 30–38 (2009).

40. Quillet, E., Labbe, L. & Queau, I. Asymmetry in sexual development of gonads in intersex rainbow trout. J. Fish Biol. 64, 1147–1151 (2004).

41. Yu, Z. H. Asymmetrical testicular weights in mammals, birds, reptiles and amphibia. Int. J. Androl. 21, 53–55 (1998).

42. Takahashi, E. et al. Migration behavior of PGCs and asymmetrical gonad formation in pond smelt Hypomesus nipponensis. Int. J. Dev. Biol. 61, 397–405 (2017).

43. Hayakawa, Y. & Kobayashi, M. Histological observations of early gonadal development to form asymmetrically in the dwarf gourami Colisa lalia. Zoolog. Sci. 29, 807–814 (2012).

44. Strüssmann, C. A. & Ito, L. S. Where does gonadal sex differentiation begin? Gradient of histological sex differentiation in the gonads of pejerrey, Odontesthes bonariensis (Pisces, Atherinidae). J. Morphol. 265, 190–196 (2005).

45. Albertson, R. C. & Yelick, P. C. Roles for fgf8 signaling in left–right patterning of the visceral organs and craniofacial skeleton. Dev. Biol. 283, 310–321 (2005).

46. Ruane, N. M., Lambert, J. G. D., Goos, H. J. T. & Komen, J. Hypocorticism and interrenal hyperplasia are not directly related to masculinization in XX mas-1/mas-1 carp, Cyprinus carpio. Gen. Comp. Endocrinol. 143, 66–74 (2005).

47. Zhai, G. et al. Characterization of Sexual Trait Development in cyp17a1-Deficient Zebrafish. Endocrinology 159, 3549–3562 (2018).

